# Coordinated HNF4α and HNF4γ Suppression Defines a Reversible Ileal Epithelial Barrier Defect in Crohn’s Disease

**DOI:** 10.1101/2025.10.27.684895

**Authors:** Debasish Halder, Amin Esmaeilniakooshkghazi, Chiranjeevi Padala, Yaohong Wang, Jason K. Hou, Xia Qiu, Lei Chen, Michael P. Verzi, Seema Khurana

## Abstract

**Background & Aims:** Ileal barrier dysfunction in Crohn’s disease (CD) can occur in macroscopically non-inflamed mucosa, persist despite clinical and endoscopic remission, and predict adverse outcomes. The epithelial mechanisms underlying this defect remain unclear. We investigated whether coordinated reduction of hepatocyte nuclear factor 4α (HNF4α) and HNF4γ defines an epithelial-intrinsic, barrier-defective state in ileal CD.

**Methods:** Barrier-associated transcriptional programs were compared in small-intestinal and colonic epithelium from wild-type mice and mice with intestinal epithelial deletion of *Hnf4α*, *Hnf4γ*, or both paralogs. Paracellular permeability was assessed in wild-type and *Hnf4α/γ* double-knockout (*Hnf4αγ*^DKO^) small-intestinal organoids. HNF4α, HNF4γ, and the direct HNF4 transcriptional target claudin-15 (CLDN15) were evaluated in human ileal CD tissue, patient-derived organoids (PDOs) generated from macroscopically non-inflamed ileal mucosa, and ileal tissue and organoids from *Tnf*^ΔARE/+^ mice. The effects of HNF4 agonists N-trans-caffeoyltyramine (NCT) and N-trans-feruloyltyramine (NFT) were assessed in CD PDOs and *Tnf*^ΔARE/+^ organoids.

**Results:** Combined, but not individual, loss of HNF4-paralog broadly disrupted small-intestinal transcriptional programs governing tight junction (TJ) organization, epithelial polarity, junctional-cytoskeletal coupling, and TJ repair, with limited effects in the colon. *Hnf4αγ*^DKO^ organoids exhibited size-selective paracellular permeability in the absence of inflammatory and stromal cells, demonstrating that combined Hnf4αγ loss is sufficient to produce an epithelial-intrinsic barrier defect. HNF4α, HNF4γ, and CLDN15 were coordinately reduced in human ileal CD epithelium and this HNF4-low phenotype was retained in PDOs from macroscopically non-inflamed ileal mucosa. Chronic TNF-driven ileitis reproduced this HNF4-low state, which was retained *ex vivo* in organoids. HNF4 agonists restored HNF4α, HNF4γ, and CLDN15 expression and improved transepithelial electrical resistance (TEER) in CD PDOs. They also increased HNF4α and CLDN15 expression in *Tnf*^ΔARE/+^ organoids.

**Conclusions:** Coordinated HNF4α/γ suppression defines an epithelial-intrinsic, barrier-defective state in ileal CD that is present in PDOs derived from macroscopically non-inflamed ileal mucosa and retained *ex vivo* without continued exposure to the inflammatory tissue environment. Its pharmacologic reversibility identifies incomplete restoration of the HNF4-dependent epithelial program as a potential mechanism of persistent ileal barrier dysfunction and supports further evaluation of HNF4-directed barrier-restorative strategies.

## Introduction

Crohn’s disease (CD) is treated predominantly as an immune-mediated inflammatory disorder, and endoscopic healing, defined by the absence of erosions or ulcerations on ileocolonoscopy, is a key therapeutic goal.^1–4^ However, increased intestinal permeability is detectable in macroscopically and histologically non-inflamed ileal mucosa, can persist despite clinical and endoscopic remission, and precedes disease onset in at-risk individuals.^5–13^ Multiple *in vivo* confocal laser endomicroscopy studies have directly visualized functional barrier defects in patients with CD and associated these abnormalities with subsequent relapse and adverse long-term disease outcomes.^5–8, 14^ The ileal localization of this permeability defect may be particularly important because ileal and ileocolonic CD are more likely than isolated colonic disease to progress from inflammatory to stricturing or penetrating behavior.^5, 15^ In the prospective ERIca trial, intestinal barrier function was assessed by confocal laser endomicroscopy in 100 patients with CD in clinical remission who were followed for a mean of 35 months.5 None of the 25 patients with terminal ileal barrier healing experienced a major adverse outcome, compared with more than 56% of patients who achieved endoscopic remission without barrier healing.5 In comparison, more than 56% of patients in endoscopic-remission only experienced a major adverse outcome.5 Terminal ileal barrier healing also predicted clinical outcomes more accurately than endoscopic or histological remission and had greater prognostic value than colonic barrier healing.^5^ Thus, ileal barrier dysfunction can persist despite conventional measures of disease control and identifies patients with CD at increased risk of adverse outcomes.^5,7, 8^

Despite its clinical and prognostic significance, intestinal permeability is not routinely assessed in CD management, and barrier healing is not an established therapeutic target.^3^ Dual-sugar tests, including the lactulose-to-mannitol ratio, can detect increased intestinal permeability but require prolonged urine collection, are affected by gastrointestinal transit and urinary excretion, lack standardized protocols, and cannot localize the barrier defect.^16–19^ Serum albumin, C-reactive protein, and commercially measured zonulin do not reliably reflect intestinal barrier function in inflammatory bowel disease (IBD).^5, 20–22^ Confocal laser endomicroscopy permits direct visualization of functional barrier defects *in vivo*, but its cost, technical complexity, and limited anatomic accessibility restrict routine use.^5^ These diagnostic limitations are compounded by an incomplete understanding of the epithelial mechanisms that sustain barrier dysfunction when conventional clinical, endoscopic, or histologic evidence of active inflammation is absent.

Mechanistic studies of intestinal barrier dysfunction have focused largely on experimental models of acute inflammatory cytokine exposure. In these models, TNF and other inflammatory cytokines activate myosin light chain kinase, reorganize the perijunctional actomyosin cytoskeleton, and induce internalization or redistribution of individual tight junction (TJ) proteins, including occludin and zonula occludens-1 (ZO-1).^23–26^ These acute cytokine-induced responses do not fully explain barrier dysfunction in non-inflamed CD mucosa or after clinical and endoscopic remission. Ultrastructural studies of CD ileum have demonstrated broader junctional abnormalities, including reduced epithelial contacts, fewer and discontinuous TJ strands, diminished junctional meshwork depth, and increased strand breaks.^4, 27, 28^ Such extensive junctional disorganization is unlikely to result solely from alteration of a single TJ protein, as illustrated by the absence of baseline intestinal permeability defects following loss of occludin or ZO-1 *in vivo*.^29, 30^ We therefore hypothesized that persistent ileal barrier dysfunction in CD reflects disruption of a coordinated epithelial transcriptional program governing junctional organization, permeability, and repair.

Experimental studies have implicated hepatocyte nuclear factor 4α (HNF4α), an epithelial-enriched nuclear receptor, in intestinal barrier regulation.^31–33^ However, intestinal epithelial *Hnf4α* knockout models have produced inconsistent effects on baseline permeability and comparatively limited changes in TJ-associated gene expression.^32–34^ These findings may reflect functional overlap between HNF4α and its paralog HNF4γ.^31^ HNF4α is broadly expressed throughout the intestinal epithelium, whereas HNF4γ is enriched in the small intestine, particularly in the distal ileum.^35, 36^ Deletion of either paralog alone produces no significant intestinal phenotype, and epithelial *Hnf4α* deletion induces compensatory *Hnf4γ* expression with minimal effects on the ileal transcriptome.^35–37^ In contrast, combined loss of *Hnf4α* and *Hnf4γ* profoundly disrupts epithelial organization and function *in vivo*.^31, 35^

Human genetic and transcriptional findings also implicate HNF4 signaling in inflammatory bowel disease (IBD). The *HNF4A* locus is associated with susceptibility to ulcerative colitis (UC) and childhood-onset CD, and reduced intestinal *HNF4A* messenger RNA (mRNA) has been reported in both diseases.^32, 38, 39^ Reduced *HNF4G* mRNA has also been reported in active UC, although its biological significance is uncertain because *HNF4G* expression is low in the normal adult colon.^40^ However, reduction of HNF4α and HNF4γ protein within the intestinal epithelium of patients with IBD has not been established, and the relationship between HNF4-paralog loss and epithelial barrier dysfunction in IBD remains unknown.

Using complementary mouse and human models, we examined whether coordinated HNF4α/γ loss disrupts epithelial barrier programs in a segment-selective manner and is sufficient to produce an epithelial-intrinsic permeability defect. Barrier-associated transcriptional programs were compared in small-intestinal and colonic epithelium following deletion of *Hnf4α*, *Hnf4γ*, or both paralogs in mice. Paracellular permeability was then assessed in s *Hnf4α/γ* double-knockout small-intestinal organoid maintained without inflammatory or stromal cells.. We examined HNF4α and HNF4γ protein expression in human ileal CD tissue and patient-derived organoids (PDOs) established from macroscopically non-inflamed ileal mucosa. Claudin-15 (CLDN15), a direct HNF4 transcriptional target, was evaluated as a representative component of the HNF4-dependent TJ program. *Tnf*^ΔARE/+^ mice and corresponding ileal organoids were used to determine whether chronic TNF-driven ileitis recapitulates the HNF4-low epithelial state. Finally, the HNF4 agonists N-trans-caffeoyltyramine (NCT) and N-trans-feruloyltyramine (NFT) were used to determine whether the associated molecular and functional abnormalities are pharmacologically reversible.

## Materials and Methods

### Human Tissue

Ileal mucosal samples were obtained from patients with active ileal Crohn’s disease (CD) and non-IBD control participants undergoing endoscopy under a protocol approved by the Institutional Review Board at Baylor College of Medicine. All participants provided written informed consent. The cohort included 19 patients with ileal CD (9 female and 10 male), including 2 patients with stricturing or penetrating disease, and 10 non-CD controls (5 female and 5 male). Macroscopically non-inflamed ileal mucosa was used for epithelial isolation and generation of patient-derived organoids (PDOs), as indicated.

### Human Patient-derived Intestinal Organoids and Morphologic Analysis

Intestinal crypts were isolated from ileal mucosal samples, embedded in Matrigel, and cultured in defined growth medium supplemented with epithelial niche factors, as previously described.^41^ Organoid were expanded before analysis, and experiments were performed between passages 3 and 8. For assessment of organoid-forming efficiency, equal numbers of crypts were plated, and the percentage that formed viable organoids by day 7 was calculated as follows: (number of viable organoids/number of plated crypts) × 100.^42^ Bright-field images of non-CD PDOs and CD PDOs were acquired under identical imaging conditions.^42^ Crypt buds were defined as epithelial protrusions extending from the central organoid body and were counted manually for each organoid.^42^ At least 50 organoids were evaluated from each independent PDO line under each condition. PDO lines from 8 patients with CD and 8 non-CD controls were analyzed, with each patient considered an independent biological replicate.

### Mouse Strains and Intestinal Organoids

*Villin-Cre^ERT2^*, *Hnf4α^f/f^*, *Hnf4γ^Crispr/Crispr^*, and *Tnf*^ΔARE/+^ mice were maintained at Rutgers University or the University of Houston under protocols approved by the respective Institutional Animal Care and Use Committees. Age- and sex-matched mice were used as previously described.^35,42^ Distal ileal tissues were collected for histologic and immunostaining analyses. Small-intestinal crypts were isolated, embedded in Matrigel, and cultured as previously described.^43^ Organoids derived from *Tnf*^+/+^ littermates served as controls for *Tnf*^ΔARE/+^ organoids.

### Pharmacologic Treatment

Matrigel-embedded organoids and organoid-derived monolayers were treated with vehicle (DMSO), 80 μmol/L *N*-trans-caffeoyltyramine (NCT, Cayman Chemical, catalog no. 40340), or 80 μmol/L *N*-trans-feruloyltyramine (NFT, MedChem Express, catalog no. HY-N2410) for 12 hours before confocal imaging or assessment of barrier function. The final DMSO concentration was identical across treatment groups.

### Organoid Permeability Assay

Small-intestinal organoids derived from wild-type (WT) and *Hnf4α/γ* double knockout (*Hnf4αγ*^DKO^) mice were exposed separately to 4-, 10-, or 70-kDa fluorescein isothiocyanate (FITC)- dextran. Luminal fluorescence accumulation was measured after 5, 30, or 60 minutes, as previously described.^44^ For each tracer and time point, luminal fluorescence was quantified in 15 randomly selected organoids from each of 3 mice per genotype. Measurements from the 15 organoids were averaged to generate one value for each mouse, which was considered the biological replicate.

### Organoid-derived epithelial monolayers and barrier assays

Human intestinal organoids were dissociated into single cells and seeded onto Transwell inserts with 0.4-μm pores (Corning) coated with type I rat-tail collagen at 20 μg/ml in 0.01% acetic acid. Barrier integrity was assessed by measuring transepithelial electrical resistance (TEER) using an EVOM2 epithelial voltohmmeter (World Precision Instruments). TEER was measured in confluent monolayers before and after 12 hours of treatment with vehicle, NCT, or NFT. Each insert was measured 3 times, and the measurements were averaged. Values were corrected for the resistance of cell-free coated inserts and multiplied by the membrane surface area to obtain Ω·cm². Measurements from replicate inserts were averaged to generate a single value per patient per treatment condition. The patient was considered the biological replicate.

### RNA Sequencing and Transcriptomic Analysis

Previously generated small-intestinal epithelial transcriptomes from WT, epithelial *Hnf4α*-knockout, *Hnf4γ*-knockout, and *Hnf4αγ*DKO mice were reanalyzed to evaluate genes involved in TJ organization, epithelial polarity, and junctional-cytoskeletal regulation.^35^ RNA sequencing was performed using colonic epithelial cells isolated from WT, epithelial *Hnf4α*-knockout, *Hnf4γ*-knockout, and *Hnf4αγ*^DKO^ mice, as previously described.^45^ RNA was extracted using the RNeasy Micro Kit (Qiagen). Transcript abundance was quantified using Kallisto version 0.44.0, and differential expression was analyzed using DESeq2 version 1.48.2 in R version 4.5.1. Bioconductor version 3.2.1. Each genotype included 3 biological replicates, with each replicate derived from an independent mouse.

### Immunohistochemistry and Immunofluorescence

Immunohistochemistry for HNF4α, HNF4γ, and claudin-15 (CLDN15) was performed on paraffin-embedded distal ileal sections from patients with ileal CD and non-CD controls, as previously described.^46^ Bright field images were acquired using an Olympus BX53 microscope under identical conditions within each staining experiment. Fluorescence images were acquired using an Olympus Fluoview FV2000 confocal microscope with a 60x objective. Fluorescence intensity was quantified using ImageJ version 1.5t. Measurements from multiple organoids or microscopic fields from the same patient or mouse were averaged before statistical analysis.

### Statistical Analysis

Statistical analyses were performed using GraphPad Prism version 11.0.2 or R version 4.5.1. Data are presented as mean ± SEM with individual biological replicate value shown where possible. The patient or mouse was considered the biological replicate. Measurements from multiple organoids, microscopic fields, replicate inserts, or repeated measurements from the same patient or mouse were treated as technical observations and averaged before statistical testing. Comparisons between 2 independent groups were performed using a 2-tailed unpaired Student *t* test for normally distributed data or a Mann–Whitney *U* test for nonnormally distributed data. Comparisons between 2 matched conditions were performed using a paired 2-tailed t test or Wilcoxon matched-pairs signed-rank test, as appropriate. Comparisons among more than 2 matched treatment conditions were performed using a repeated-measures analysis of variance followed by a multiplicity-adjusted post hoc test or a Friedman test followed by Dunn’s multiple-comparison test, as appropriate. The statistical test and biological sample size for each experiment are reported in the corresponding figure legend. All tests were 2-sided, and *P*<.05 was considered statistically significant.

## Results

### Combined loss of HNF4α and HNF4γ selectively disrupts epithelial barrier programs in the small intestine

To determine whether HNF4 paralogs coordinately regulate intestinal epithelial barrier programs, we performed a barrier-focused reanalysis of our published small-intestinal epithelial transcriptomes from wild-type (WT), *Hnf4α*-knockout (*Hnf4α*^KO^), Hnf4γ-knockout (*Hnf4γ*^KO^), and *Hnf4α/γ* double-knockout (*Hnf4αγ*^DKO^) mice.^35^ Although these transcriptomes were originally generated to investigate HNF4-dependent enterocyte differentiation, their implications for epithelial barrier regulation had not been systematically examined.

Combined deletion of *Hnf4α* and *Hnf4γ* broadly disrupted transcriptional programs governing TJ assembly and function, epithelial polarity, and junctional-cytoskeletal organization. In contrast, deletion of either paralog alone produced no significant changes in these transcriptional programs, consistent with functional compensation between HNF4α and HNF4γ. Genes reduced in *Hnf4αγ*^DKO^ epithelium included components of TJ assembly and scaffolding (*F11r*, *Magi3*, *Tjap1*, and *Marveld3*), apical junctional organization and epithelial polarity (*Tjp2*, *Pard6b*, and *Prkcz*), and junctional-cytoskeletal coupling (*Myo1e*). Multiple regulators of RhoA-dependent cytoskeletal dynamics and TJ repair, including members of the *Arhgef*, *Arhgap*, and *Pak* families, were also dysregulated.^35^ Barrier-associated claudins *Cldn4* and *Cldn7* and the small-intestine-enriched *Cldn15* were reduced, whereas the pore-forming claudin *Cldn2* was increased. Analysis of published ileal CD transcriptomic datasets identified a similar expression pattern, with reduced *CLDN4*, *CLDN7*, *CLDN15*, *MAGI3*, *F11R*, and *TJP2* expression and increased *CLDN2*.^47–49^

To determine whether the requirement for combined HNF4-paralog activity differed between intestinal segments, we performed RNA sequencing of colonic epithelium from the same four mouse genotypes and analyzed genes associated with TJ organization, epithelial polarity, and cytoskeletal regulation. In contrast to the broad disruption of the small-intestinal barrier program, only five TJ or TJ-associated genes were significantly altered in *Hnf4αγ*^DKO^ colonic epithelium at an adjusted *P* value < .01 (Table 1), and most changes were modest. *Cldn23, Ocln, Cldn3,* and *Prkcz* were downregulated, whereas *Cldn2* was upregulated; only *Cldn23* changed by more than twofold. Thus, the transcriptional consequences of combined HNF4α/γ loss were strongly segment selective, with broad disruption of barrier-associated programs in the small intestine and comparatively limited effects in the colon.

**Table 1.** Differential gene expression in *Hnf4αγ*^DKO^ colonic epithelium. RNA sequencing was performed on colonic epithelial cells isolated from *Hnf4αγ*^DKO^ mice and wild-type (WT) controls, with three independent mice per genotype. Transcript abundance was quantified using Kallisto, and differential expression was analyzed using DESeq2. Positive log2 fold-change values indicate increased expression in *Hnf4αγ*^DKO^ relative to WT epithelium, whereas negative values indicate decreased expression. The workbook provides the complete differential-expression results and filtered results based on a Benjamini–Hochberg-adjusted *P* value < .01. Genes with log2 fold change > 1 or < −1 were classified as upregulated or downregulated, respectively. Reported variables include Ensembl gene identifier, mean normalized expression across samples (baseMean), log2 fold change, standard error of the log2 fold change (lfcSE), Wald statistic, raw *P* value, adjusted *P* value, −log10-adjusted *P* value, and MGI gene symbol. Among TJ and TJ-associated genes, *Cldn23, Ocln, Cldn3,* and *Prkcz* were significantly downregulated and *Cldn2* was significantly upregulated at adjusted *P* < .01; only *Cldn23* had an absolute log2 fold change > 1.

### HNF4-paralog loss causes size-selective paracellular permeability

To determine whether the transcriptional changes produced by combined HNF4-paralog loss were accompanied by functional barrier impairment, we assessed molecular size-dependent paracellular permeability in three-dimensional WT and *Hnf4αγ*^DKO^ small-intestinal organoids using fluorescent dextran tracers.^50^ Organoids were exposed separately to 4-, 10-, or 70-kDa FITC-dextran, and luminal tracer accumulation was measured over time.^44, 50–52^ WT organoids excluded all 3 tracers from the lumen for up to 30 minutes, consistent with an intact epithelial barrier (Figure 1). In contrast, *Hnf4αγ*^DKO^ organoids exhibited luminal accumulation of the 4- and 10-kDa tracers at 30 minutes while continuing to exclude the 70-kDa tracer. This pattern demonstrated size-selective paracellular permeability without gross loss of epithelial integrity. Thus, combined HNF4α/γ loss was sufficient to produce an epithelial-intrinsic barrier defect in organoids maintained without inflammatory or stromal cells.

**Figure 1.**
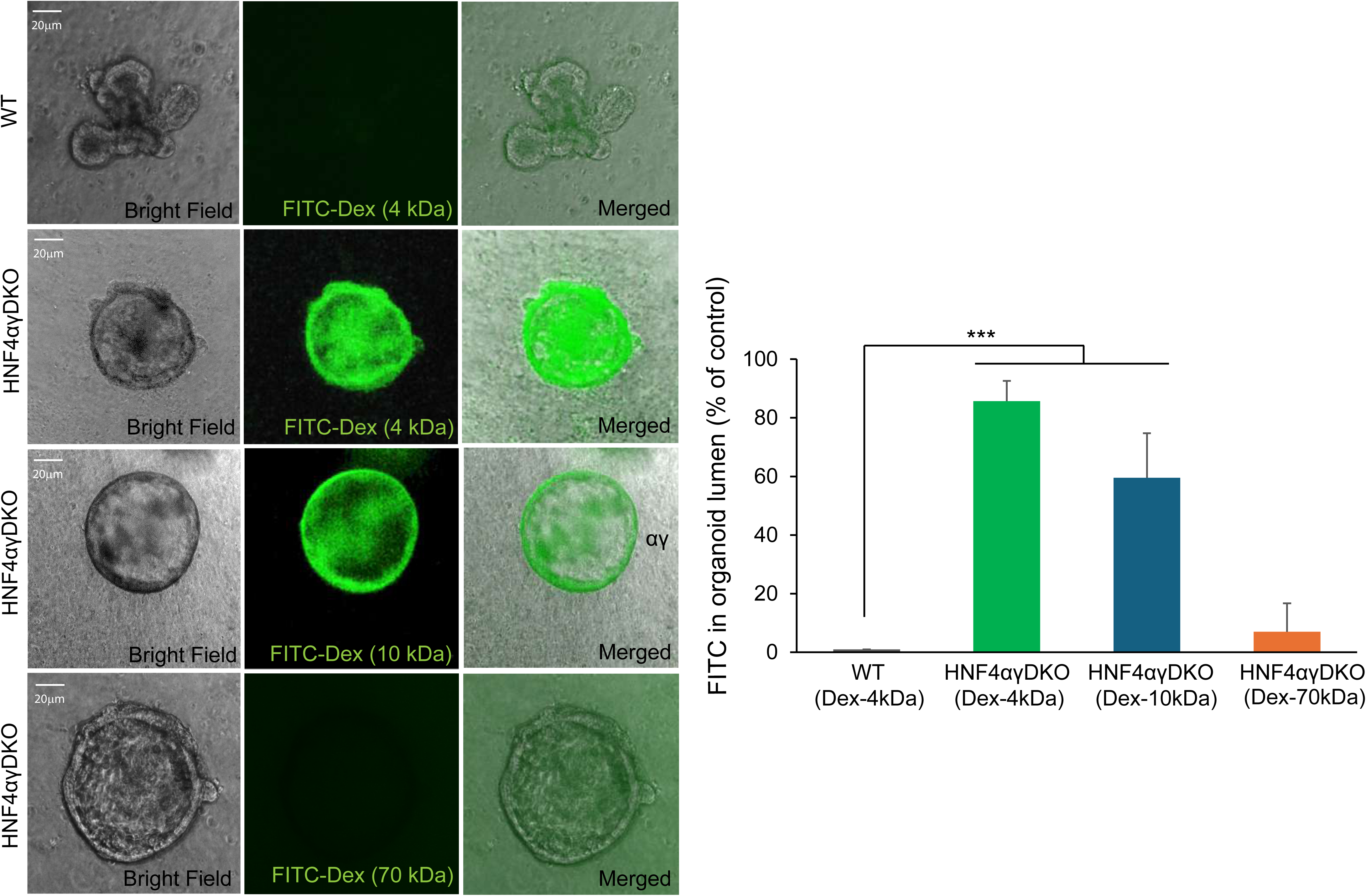
Combined loss of HNF4α and HNF4γ causes size-selective paracellular permeability in small-intestinal organoids. **(A)** Representative bright-field, FITC-fluorescence, and merged images of small-intestinal organoids derived from WT and *Hnf4αγ*^DKO^ mice after 30 minutes of exposure to 4-, 10-, or 70-kDa FITC-dextran. WT organoids excluded 4-kDa FITC-dextran from the lumen, whereas *Hnf4αγ*^DKO^ organoids accumulated 4- and 10-kDa FITC-dextran. The 70-kDa tracer remained excluded from *Hnf4αγ*^DKO^ organoids, demonstrating size-selective permeability without gross epithelial disruption. Organoids were cultured in medium supplemented with 25 mmol/L acetate to compensate for the acetyl-CoA deficiency in *Hnf4αγ*^DKO^ organoids.^43^ **(B)** Quantification of luminal FITC-dextran accumulation after 30 minutes. Data are presented as mean ± SEM from three independent mice per genotype. Fifteen organoids were analyzed per mouse and condition and averaged to generate each biological-replicate value. Statistical significance between WT and *Hnf4αγ*^DKO^ organoids exposed to FITC-dextran was determined using a two-tailed unpaired Student *t* test. \*\*\**P* < .001. Scale bars, 20 μm.

### HNF4α and HNF4γ are coordinately reduced in ileal CD epithelium

We next determined whether HNF4α and HNF4γ protein expression was reduced in human ileal CD epithelium. CLDN15 was examined as a representative HNF4-dependent TJ target protein because HNF4α directly binds the *Cldn15* promoter and regulates its transcription.^33^ Moreover, *Cldn15* expression was significantly reduced in *Hnf4αγ*^DKO^ small-intestinal epithelium, and reduced *CLDN15* expression has been reported in terminal ileal biopsies from patients with active CD.^49^ HNF4α, HNF4γ, and CLDN15 were examined by immunohistochemistry in distal ileal biopsies from patients with ileal CD and non-CD controls. In non-CD ileal epithelium, HNF4α, detected using an antibody that recognizes both P1- and P2-derived isoforms, showed strong nuclear staining along the crypt-villus axis (Figure 2). HNF4γ showed nuclear and cytoplasmic staining, whereas CLDN15 was strongly expressed throughout the epithelium. In contrast, ileal CD epithelium showed coordinated reductions in nuclear HNF4α, epithelial HNF4γ and CLDN15 staining within the same tissue samples (Figure 2). These findings provide protein-level evidence of coordinated reductions in HNF4-paralog in ileal CD epithelium, accompanied by reduced expression of HNF4-dependent TJ component.

**Figure 2.**
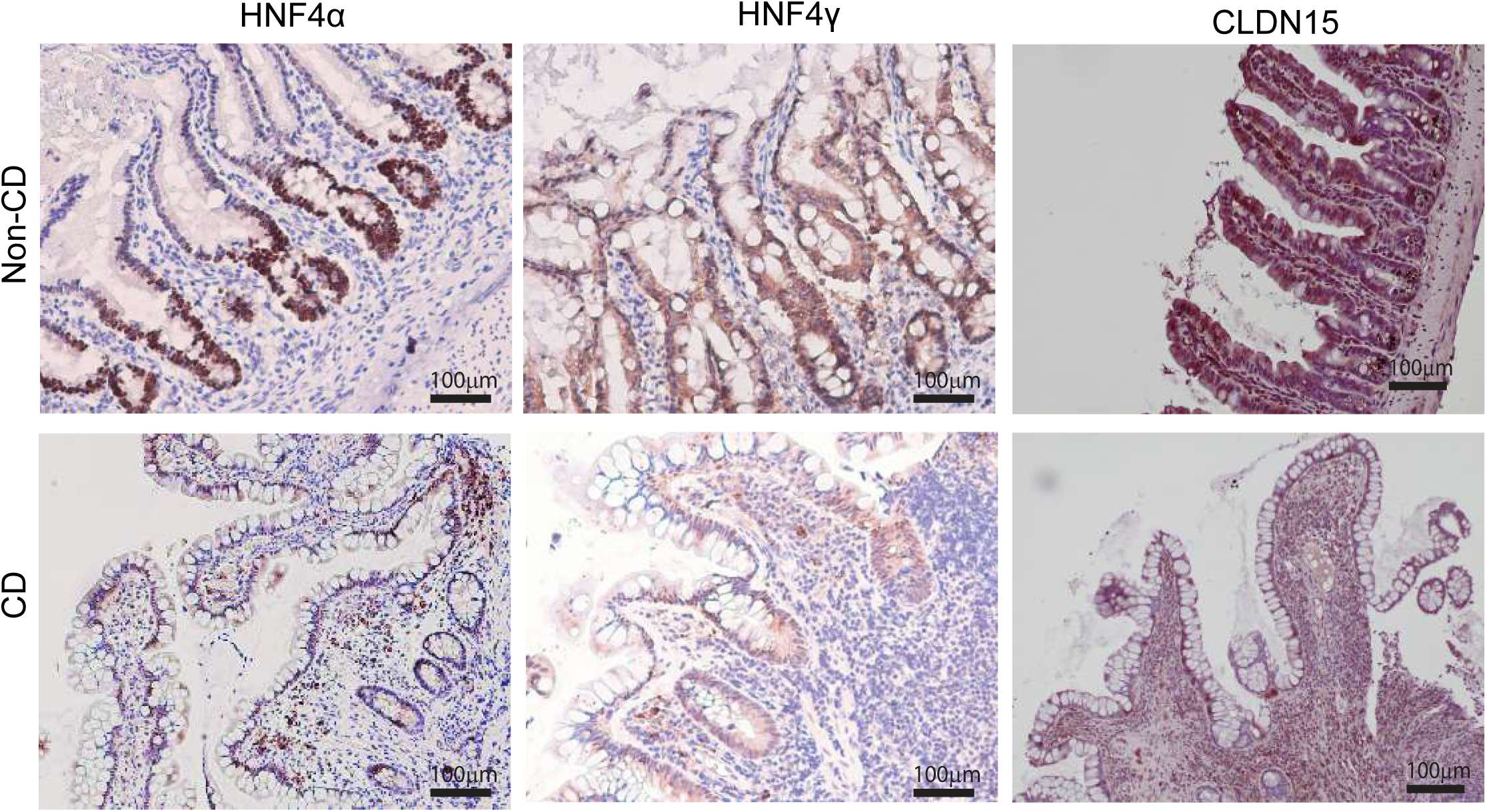
Coordinated reduction of HNF4α, HNF4γ, and CLDN15 in distal ileal epithelium from patients with Crohn’s disease. Representative immunohistochemical staining of HNF4α, HNF4γ, and CLDN15 in paraffin-embedded distal ileal sections from non-CD controls and patients with ileal Crohn’s disease (CD). In non-CD ileal epithelium, HNF4α showed strong nuclear localization along the crypt-villus axis, HNF4γ was detected in nuclear and cytoplasmic compartments, and CLDN15 showed strong epithelial staining. Ileal epithelium from patients with CD showed coordinated reductions in nuclear HNF4α, HNF4γ, and CLDN15 staining. Images are representative of 11 patients with CD and 4 non-CD controls. Scale bars, 100 μm.

### The HNF4-low phenotype is retained in CD organoids derived from macroscopically non-inflamed ileal mucosa

To determine whether HNF4-paralog suppression was retained in epithelial cultures derived from macroscopically non-inflamed CD mucosa, we examined patient-derived organoids (PDOs) established from distal ileum of patients with active ileal CD and non-CD controls. HNF4α protein expression was significantly reduced in CD PDOs compared with non-CD PDOs (*P*<.0001; n=8 patients per group; Figure 3A). CLDN15 expression was reduced in parallel, consistent with diminished HNF4-dependent transcriptional activity (Figure 3A).^33^ HNF4γ was also markedly reduced in the same CD PDOs and was nearly undetectable compared with its robust expression in non-CD controls (*P*<.0001; n=8 patients per group; Figure 3B). Because these PDOs were derived from macroscopically non-inflamed mucosa and maintained without immune or stromal cells, retention of coordinated HNF4α, HNF4γ, and CLDN15 reduction *ex vivo* supports an epithelial-intrinsic state that does not require continued exposure to the inflammatory tissue environment. This interpretation is consistent with previous evidence that ileal CD PDOs retain disease-associated epithelial transcriptional programs *ex vivo*.^53^

**Figure 3A.**
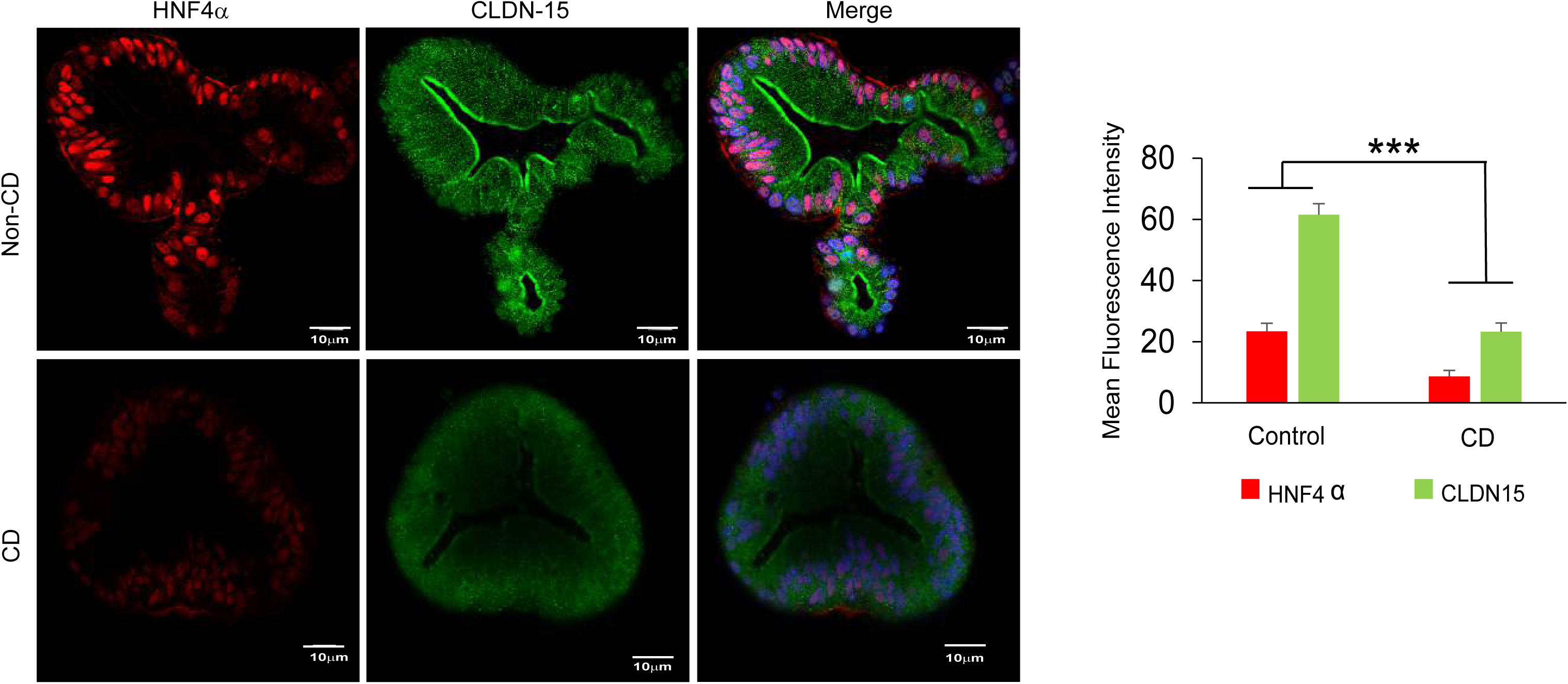
Reduced HNF4α and CLDN15 expression in Crohn’s disease patient-derived organoids. Representative confocal immunofluorescence images of patient-derived organoids (PDOs) generated from macroscopically non-inflamed distal ileal mucosa of patients with ileal Crohn’s disease (CD) and non-CD controls. Organoids were stained for HNF4α (red) and CLDN15 (green), and nuclei were counterstained with DAPI (blue). Non-CD PDOs showed strong nuclear HNF4α and prominent CLDN15 localization along the apical epithelial membrane. CD PDOs showed markedly reduced HNF4α and CLDN15 fluorescence. Right panels show quantification of HNF4α and CLDN15 mean fluorescence intensities. Data are presented as mean ± SEM from PDOs derived from 8 patients per group. Statistical significance was determined using a two-tailed unpaired Student *t* test. ***P<0.001.

**Figure 3B.**
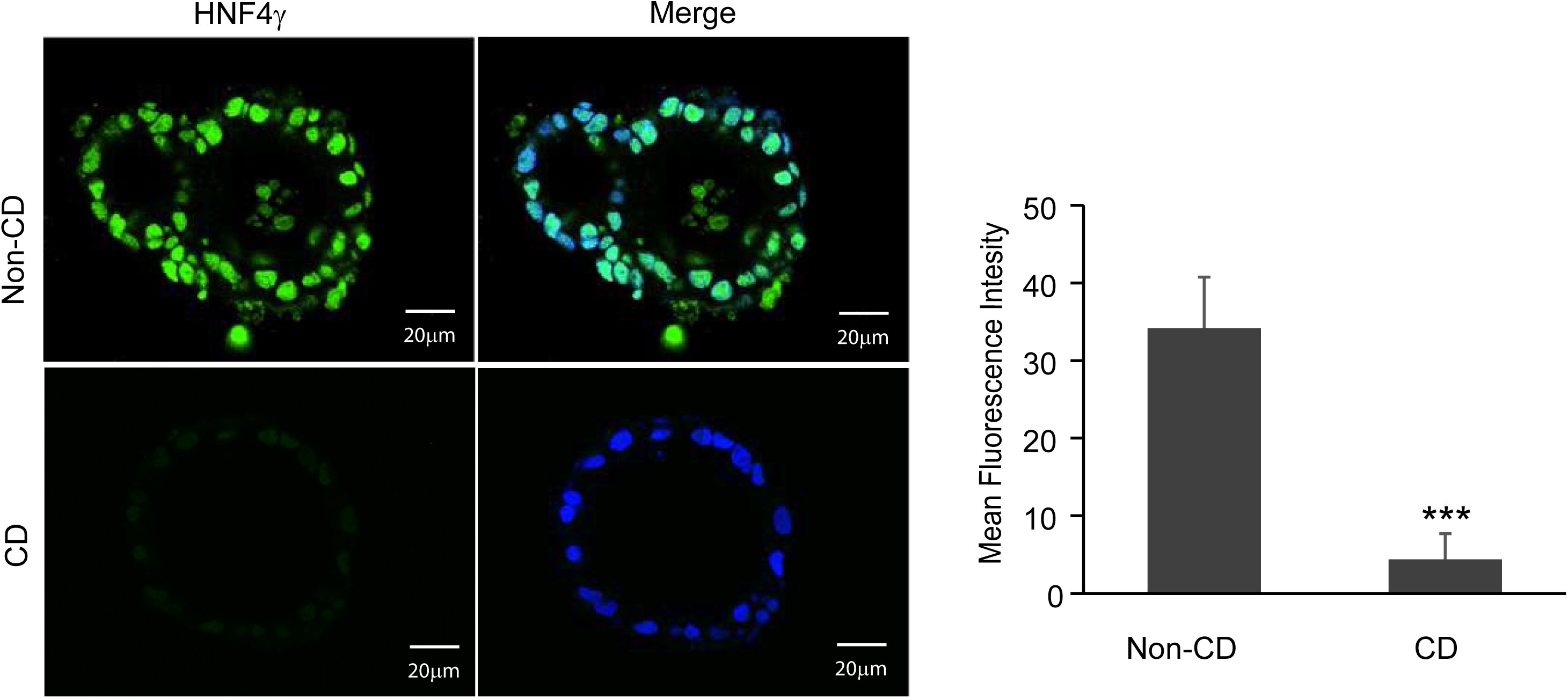
Reduced HNF4γ expression in Crohn’s disease patient-derived organoids. Representative confocal immunofluorescence images of patient-derived organoids (PDOs) generated from macroscopically non-inflamed distal ileal mucosa of patients with ileal Crohn’s disease (CD) and from non-CD controls. Organoids were stained for HNF4γ (green), and nuclei were counterstained with DAPI (blue). Non-CD PDOs showed abundant epithelial HNF4γ expression, whereas CD PDOs showed markedly reduced HNF4γ fluorescence. Right panel shows quantification of HNF4γ mean fluorescence intensity. Data are presented as mean ± SEM from PDOs derived from 8 patients per group. Statistical significance was determined using a two-tailed unpaired Student *t* test. ***P<0.001. Scale bar, 20 μm.

### Chronic TNF-driven ileitis recapitulates HNF4-paralog suppression

To determine whether chronic ileitis is associated with suppression of HNF4 paralogs *in vivo*, we examined distal ileal tissue from *Tnf*^ΔARE/+^ mice, which develop spontaneous chronic TNF-driven ileitis, and *Tnf*^+/+^ controls. In control ileum, HNF4α showed strong nuclear localization along the crypt–villus axis, whereas HNF4γ was detected throughout the epithelium (Figure 4A). In contrast, inflamed distal ileum from *Tnf*^ΔARE/+^ mice showed markedly reduced nuclear HNF4α and epithelial HNF4γ staining (Figure 4A; n=6 mice per genotype). Thus, chronic TNF-driven ileitis was associated with coordinated reduction of both HNF4 paralogs in the ileal epithelium *in vivo*. We next determined whether this HNF4-low phenotype was retained after removal from the inflammatory tissue environment. HNF4α and CLDN15 were significantly reduced in organoids established from *Tnf*^ΔARE/+^ distal ileum compared with *Tnf*^+/+^ controls (*P*<.0001; n=6 mice per genotype; Figure 4B). HNF4γ was also markedly reduced in *Tnf*^ΔARE/+^ organoids (*P*<.0001; n=6 mice per genotype; Figure 4C). Thus, coordinated reduction of HNF4α, HNF4γ, and CLDN15 was retained *ex vivo* without continued exposure to the inflammatory tissue microenvironment. This phenotype paralleled that observed in human CD PDOs derived from macroscopically non-inflamed ileal mucosa.

**Figure 4A.**
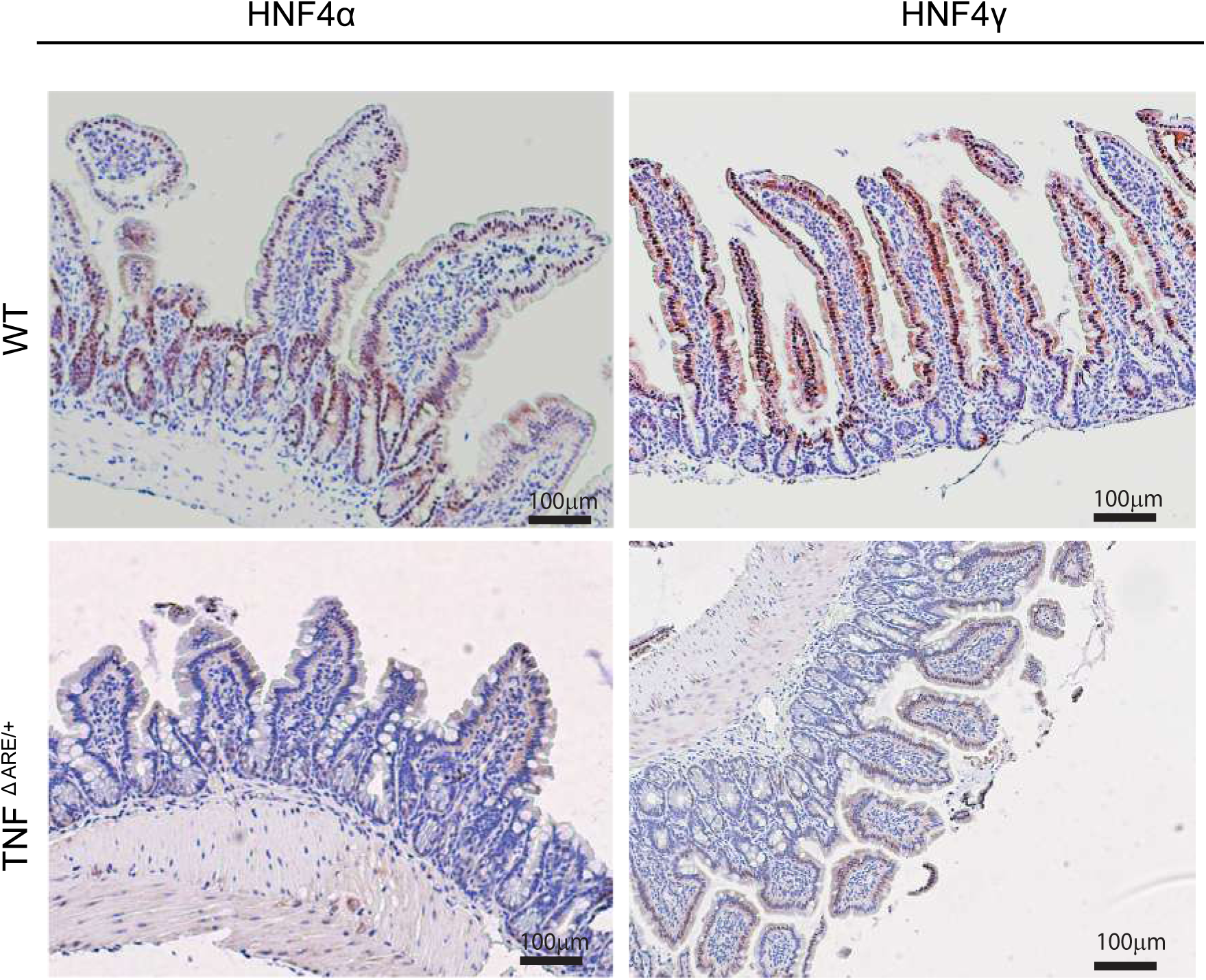
Reduced epithelial HNF4α and HNF4γ expression in chronic TNF-driven ileitis. Representative immunohistochemical staining of HNF4α and HNF4γ in paraffin-embedded distal ileal sections from *Tnf*^+/+^ and *Tnf*^ΔARE/+^ mice. In *Tnf*^+/+^ controls, HNF4α showed strong nuclear localization, whereas HNF4γ showed abundant epithelial staining along the crypt–villus axis. In contrast, inflamed ileal epithelium from *Tnf*^ΔARE/+^ mice showed markedly reduced HNF4α and HNF4γ staining. Images are representative of 6 mice per genotype. Scale bars, 100 μm.

**Figure 4B.**
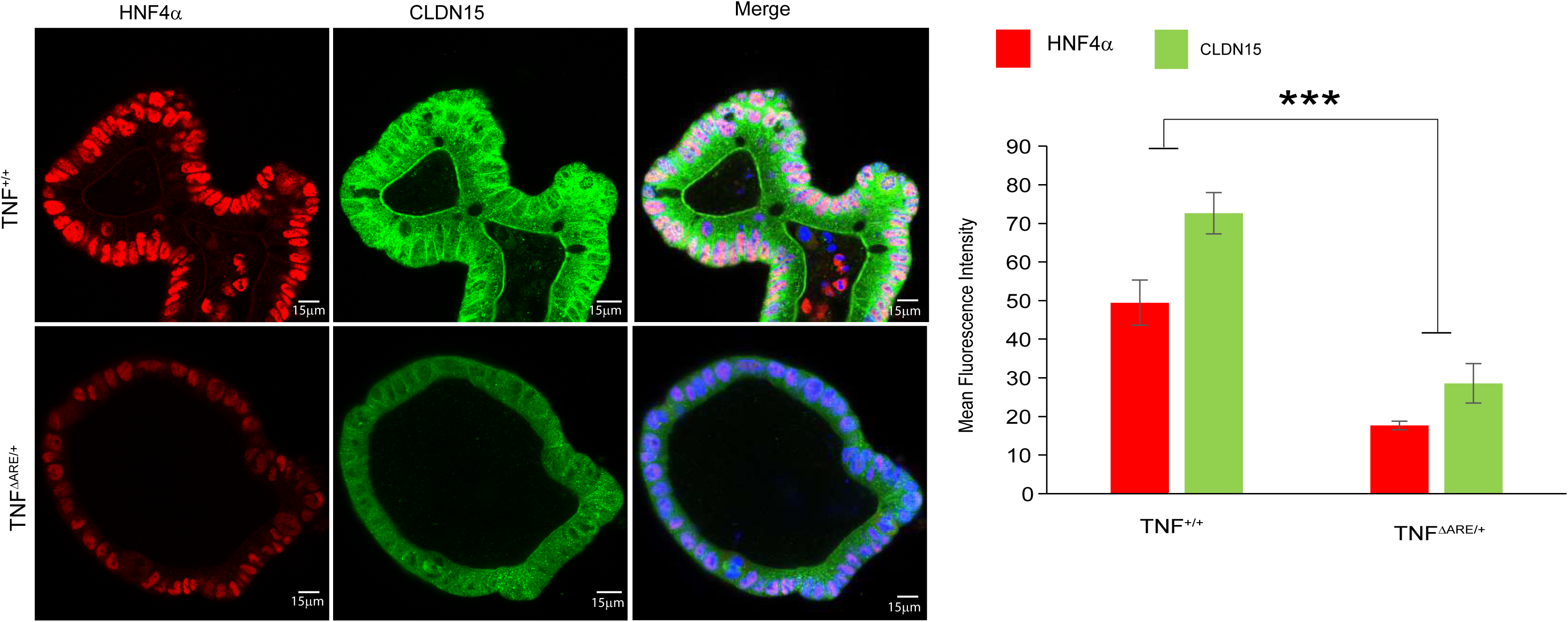
Reduced HNF4α and CLDN15 expression in ileal organoids derived from *Tnf*^ΔARE/+^ mice. Representative confocal immunofluorescence images of organoids derived from the distal ileum of *Tnf*^+/+^ and *Tnf*^ΔARE/+^ mice. Organoids were stained for HNF4α (red) and CLDN15 (green), and nuclei were counterstained with DAPI (blue). *Tnf*^+/+^ organoids showed strong nuclear HNF4α and prominent CLDN15 localization along the apical epithelial membrane, whereas *Tnf*^ΔARE/+^ organoids showed markedly reduced HNF4α and CLDN15 fluorescence. Although the anti-human CLDN15 antibody produced nonspecific background fluorescence in mouse organoids, apical membrane-associated CLDN15 staining was reduced in *Tnf*^ΔARE/+^ organoids. Right panels show quantification of HNF4α and CLDN15 mean fluorescence intensity. Data are presented as mean ± SEM from 6 mice per genotype. Statistical significance was determined using a two-tailed unpaired Student *t* test. ****P<0.001. Scale bar, 15 μm.

**Figure 4C.**
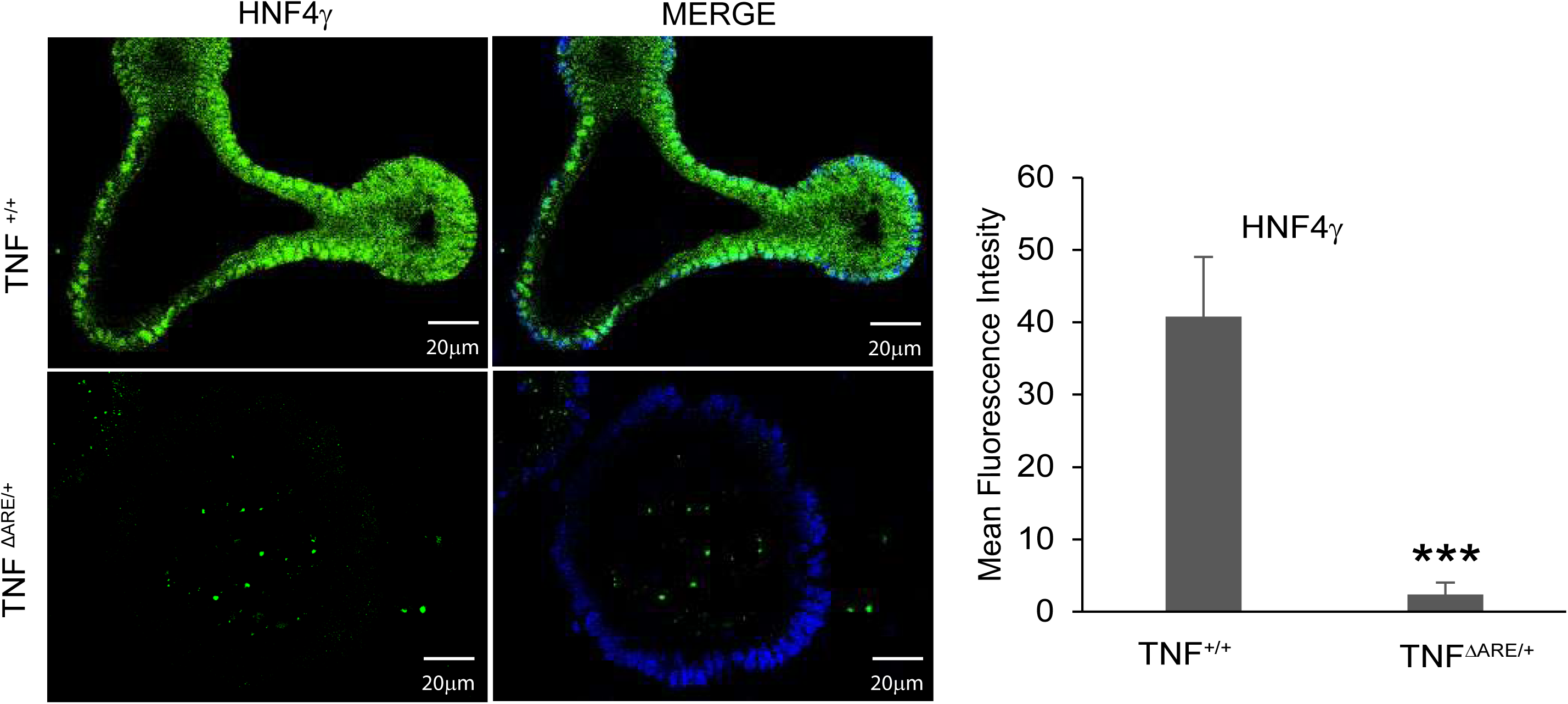
Reduced HNF4γ expression in ileal organoids derived from *Tnf*^ΔARE/+^ mice. Representative confocal immunofluorescence images of organoids derived from the distal ileum of *Tnf*^+/+^ and *Tnf*^ΔARE/+^ mice. Organoids were stained for HNF4γ (green), and nuclei were counterstained with DAPI (blue). *Tnf*^+/+^ organoids showed abundant epithelial HNF4γ expression, whereas *Tnf*^ΔARE/+^ organoids showed markedly reduced HNF4γ fluorescence. Right panel shows quantification of HNF4γ mean fluorescence intensity. Data are presented as mean ± SEM from 6 mice per genotype. Statistical significance was determined using a two-tailed unpaired Student *t* test. ***P<0.001. Scale bar, 20 μm.

### NCT and NFT reverse the HNF4-low molecular phenotype in human CD and chronic *Tnf*^ΔARE/+^ organoids

We next tested whether the HNF4-low epithelial phenotype could be reversed pharmacologically. N-trans-caffeoyltyramine (NCT) and N-trans-feruloyltyramine (NFT) are naturally occurring hydroxycinnamic acid amides identified as HNF4α agonists through an HNF4α-responsive transcriptional screen.^54^ Both compounds activate HNF4α-dependent transcription, and HNF4α knockdown abolishes their activity.^54^ Drug-affinity responsive target stability assays further support direct interaction with HNF4α, whereas neither compound exhibits estrogen-receptor or PPARγ agonist activity.^54^ CD PDOs were treated with NCT or NFT at 80 μmol/L for 12 hours. Both compounds significantly increased HNF4α and CLDN15 expression toward levels observed in non-CD controls (P<.0001; n=8 patients per group; Figure 5A). NCT and NFT also increased HNF4γ expression in CD PDOs (Figure 5B). Recovery of CLDN15, a direct HNF4α transcriptional target, was consistent with increased activation of HNF4-dependent transcription. To determine whether this response was reproduced in chronic TNF-driven ileitis, organoids derived from *Tnf*^ΔARE/+^ distal ileum were treated with NCT or NFT under the same conditions. Both compounds increased HNF4α and CLDN15 expression toward levels observed in *Tnf*^+/+^ control organoids (*P*<.0001; n=6 mice per genotype; Figure 5C). Thus, the HNF4-low molecular phenotype was pharmacologically reversible in human CD PDOs and organoids derived from mice with chronic TNF-driven ileitis.

**Figure 5A.**
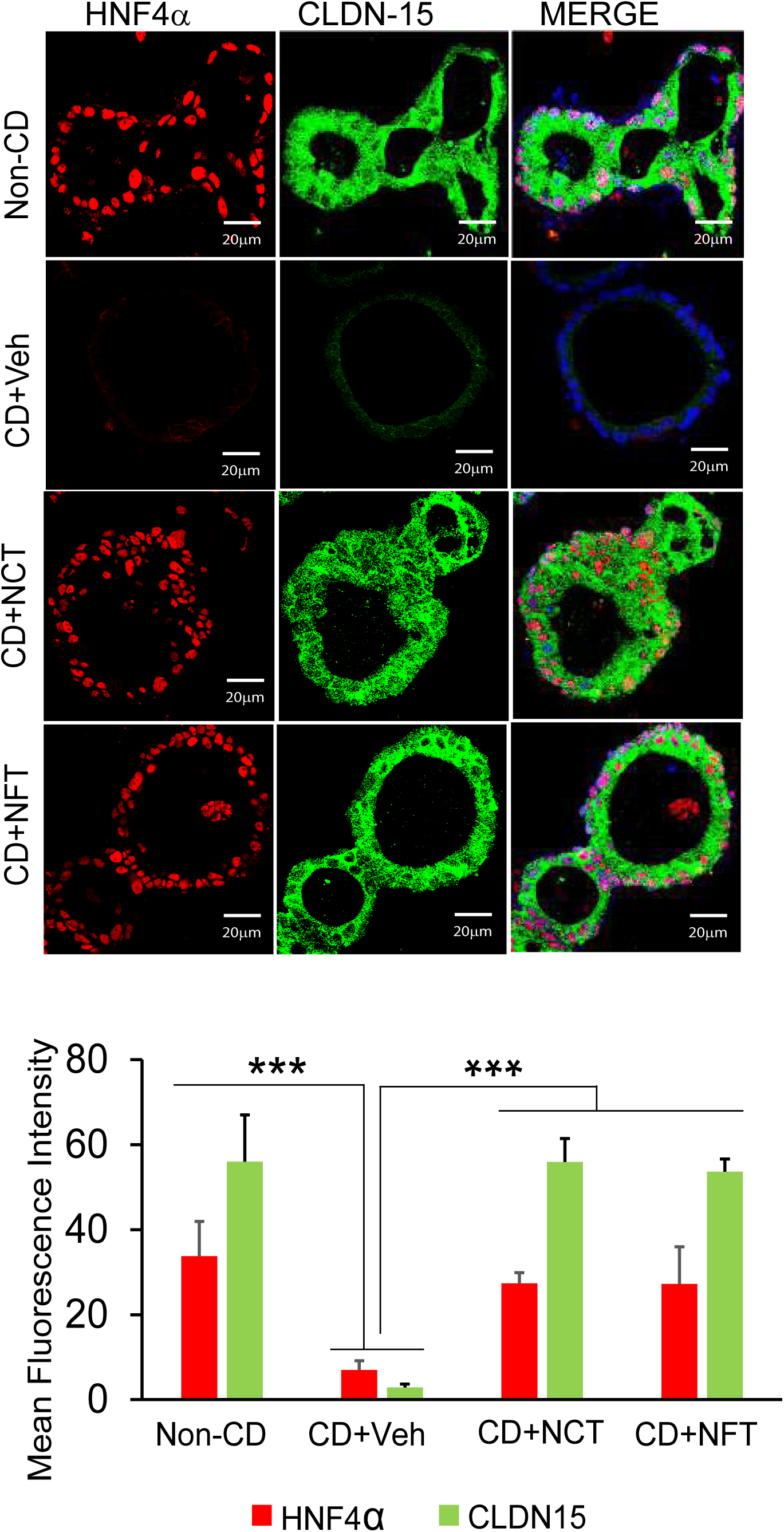
HNF4 agonists restore HNF4α and CLDN15 expression in Crohn’s disease patient-derived organoids. Representative confocal immunofluorescence images of patient-derived organoids (PDOs) generated from macroscopically non-inflamed distal ileal mucosa of patients with Crohn’s disease (CD) and from non-CD controls. Non-CD PDOs were treated with vehicle (Veh; DMSO), N-trans-caffeoyltyramine (NCT; 80 μmol/L), or N-trans-feruloyltyramine (NFT; 80 μmol/L) for 12 hours. PDOs were stained for HNF4α (red) and CLDN15 (green), and nuclei were counterstained with DAPI (blue). Vehicle-treated non-CD PDOs showed strong nuclear HNF4α and apical CLDN15 localization, whereas vehicle-treated CD PDOs showed markedly reduced HNF4α and CLDN15 fluorescence. NCT and NFT restored HNF4α and CLDN15 expression in CD PDOs toward non-CD levels. Bottom panels show quantification of HNF4α and CLDN15 mean fluorescence intensities. Data are presented as mean ± SEM from PDOs derived from 8 patients per group. Non-CD versus CD comparisons were performed using two-tailed unpaired Student *t* tests; paired comparisons of vehicle- and agonist-treated CD PDOs were performed using two-tailed paired Student *t* tests. ***P < .001. Scale bars, 20 μm.

**Figure 5B.**
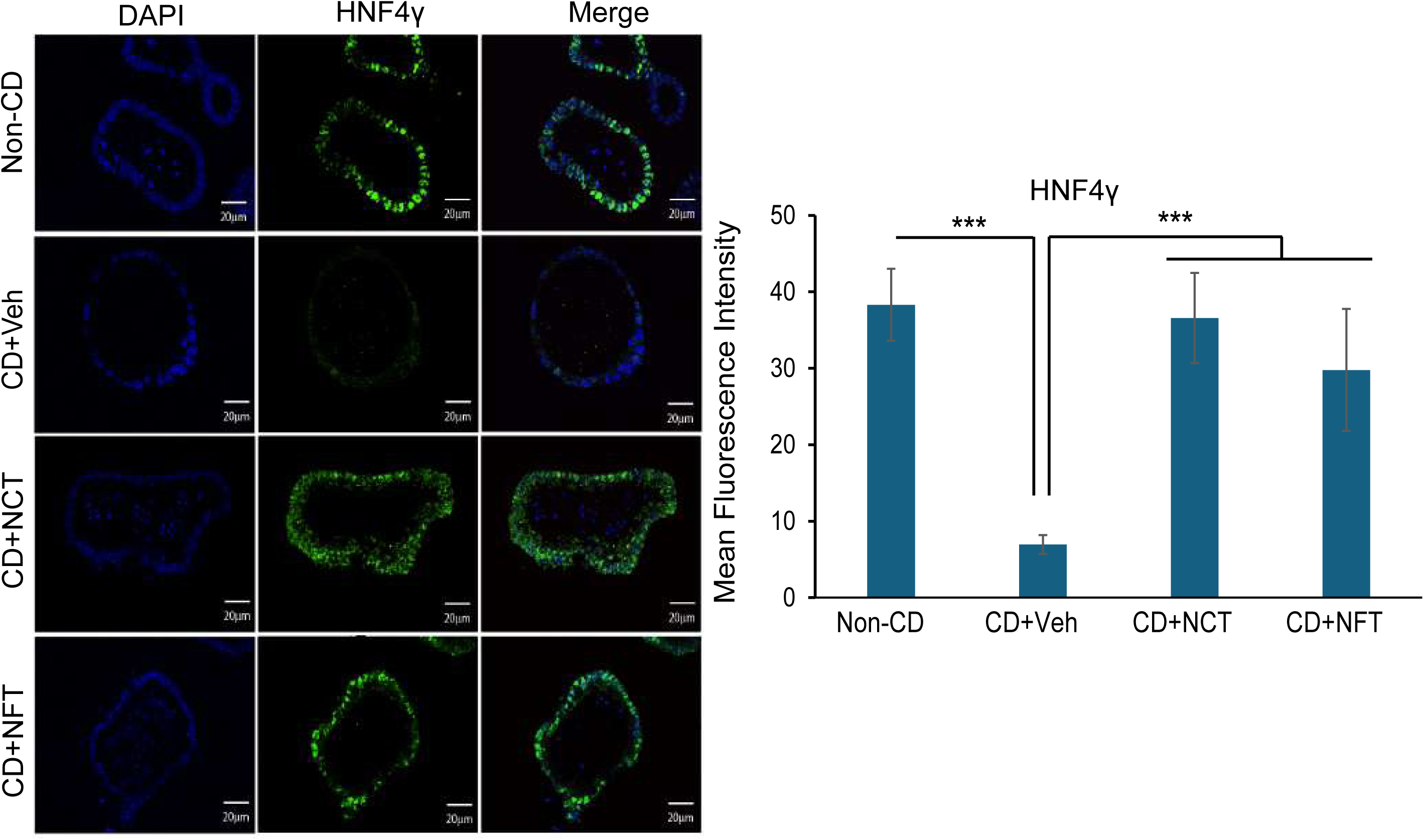
HNF4 agonists restore HNF4γ expression in Crohn’s disease patient-derived organoids. Representative confocal immunofluorescence images of patient-derived organoids (PDOs) generated from macroscopically non-inflamed distal ileal mucosa of patients with Crohn’s disease (CD) and non-CD controls. Non-CD PDOs were treated with vehicle, and CD PDOs were treated with vehicle (Veh; DMSO), *N*-trans-caffeoyltyramine (NCT; 80 μmol/L), or *N*-trans-feruloyltyramine (NFT; 80 μmol/L) for 12 hours. PDOs were stained for HNF4γ (green), and nuclei were counterstained with DAPI (blue). Vehicle-treated non-CD PDOs showed abundant epithelial HNF4γ staining, whereas vehicle-treated CD PDOs showed markedly reduced HNF4γ fluorescence. NCT and NFT restored HNF4γ expression in CD PDOs toward non-CD levels. Right, quantification of HNF4γ mean fluorescence intensity. Data are presented as mean ± SEM from PDOs derived from 8 non-CD controls and 8 patients with CD. The non-CD versus CD comparison was performed using a two-tailed unpaired Student *t* test; paired comparisons of vehicle- and agonist-treated CD PDOs were performed using two-tailed paired Student *t* tests. ***P < .001. Scale bars, 20 μm.

**Figure 5C.**
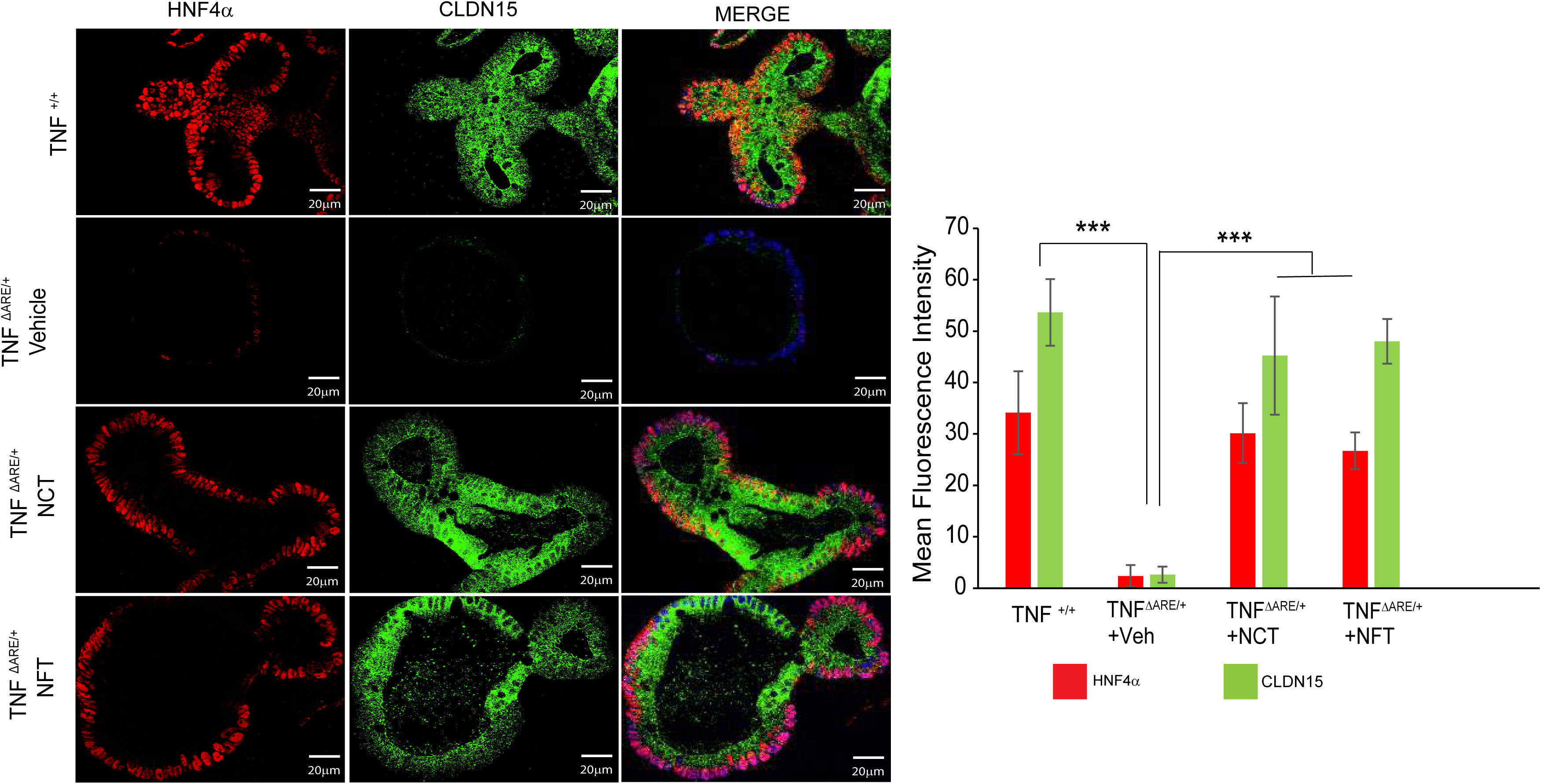
HNF4 agonists restore HNF4α and CLDN15 expression in ileal organoids derived from *Tnf*^ΔARE/+^ mice. Representative confocal immunofluorescence images of organoids derived from the distal ileum of *Tnf*^+/+^ and *Tnf*^ΔARE/+^ mice. *Tnf*^+/+^ organoids were treated with vehicle, and *Tnf*^ΔARE/+^ organoids were treated with vehicle (Veh; DMSO), *N*-trans-caffeoyltyramine (NCT; 80 μmol/L), or *N*-trans-feruloyltyramine (NFT; 80 μmol/L) for 12 hours. Organoids were stained for HNF4α (red) and CLDN15 (green), and nuclei were counterstained with DAPI (blue). Vehicle-treated *Tnf*^+/+^ organoids showed strong nuclear HNF4α staining and apical CLDN15 localization, whereas vehicle-treated *Tnf*^ΔARE/+^ organoids showed markedly reduced HNF4α and CLDN15 fluorescence. NCT and NFT restored HNF4α and CLDN15 expression in *Tnf*^ΔARE/+^ organoids toward *Tnf*^+/+^ levels. Right, quantification of HNF4α and CLDN15 mean fluorescence intensities. Data are presented as mean ± SEM from organoids derived from 6 mice per genotype. The *Tnf*^+/+^ versus *Tnf*^ΔARE/+^ vehicle comparison was performed using two-tailed unpaired Student *t* tests; paired comparisons of vehicle- and agonist-treated *Tnf*^ΔARE/+^ organoids were performed using two-tailed paired Student *t* tests. Scale bars, 20 μm.

### NCT and NFT improve epithelial barrier function in CD PDO monolayers

Finally, we determined whether recovery of HNF4α, HNF4γ, and CLDN15 expression was accompanied by improved epithelial barrier function. CD and non-CD organoids were dissociated and cultured as confluent epithelial monolayers on permeable supports. Vehicle-treated CD PDO monolayers exhibited significantly lower TEER than vehicle treated non-CD monolayers (P < .0001; n = 8 patients per group; Figure 6). Treatment with NCT or NFT significantly increased TEER in CD monolayers toward non-CD levels. Absolute TEER values were within previously reported ranges for human ileal organoid-derived monolayers, which vary according to differentiation protocol, culture conditions, and Transwell configuration.^55–57^ Accordingly, concurrently cultured non-CD monolayers provided the most appropriate benchmark for treatment response. These findings demonstrate that NCT and NFT improved epithelial electrical resistance in CD patient-derived monolayers.

**Figure 6.**
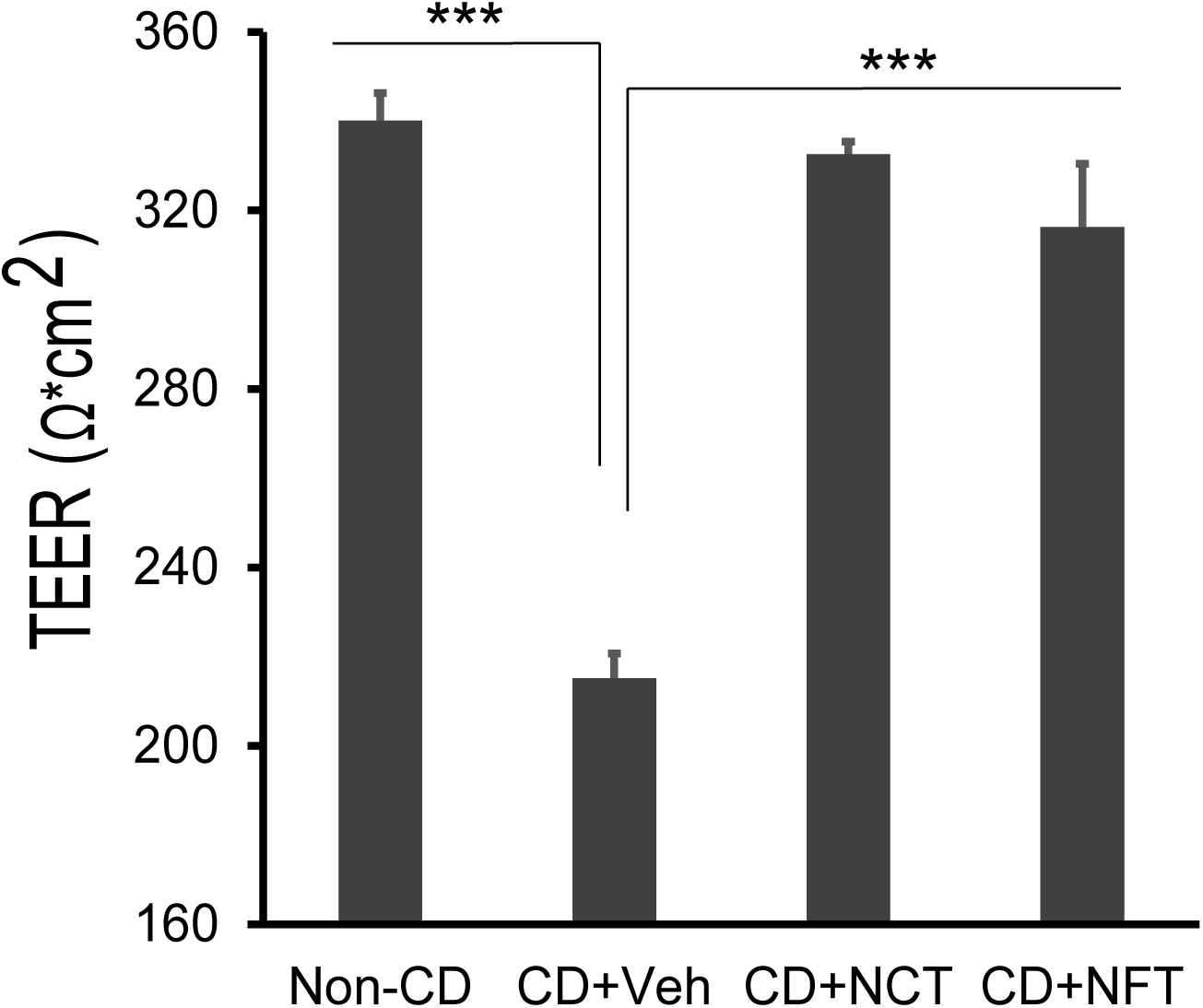
HNF4 agonists restore transepithelial electrical resistance in Crohn’s disease patient-derived organoid monolayers. Transepithelial electrical resistance (TEER) was measured in patient-derived organoid (PDO) monolayers generated from macroscopically non-inflamed distal ileal mucosa of patients with Crohn’s disease (CD) and non-CD controls. Non-CD PDO monolayers were treated with vehicle, and CD PDO monolayers were treated with vehicle (Veh; DMSO), *N*-trans-caffeoyltyramine (NCT; 80 μmol/L), or *N*-trans-feruloyltyramine (NFT; 80 μmol/L) for 12 hours. Vehicle-treated CD PDO monolayers showed reduced TEER compared with vehicle-treated non-CD monolayers. NCT and NFT increased TEER in CD PDO monolayers toward non-CD levels. Data are presented as mean ± SEM from PDOs derived from 8 non-CD controls and 8 patients with CD. The non-CD versus CD comparison was performed using a two-tailed unpaired Student *t* test; paired comparisons of vehicle- and agonist-treated CD PDO monolayers were performed using two-tailed paired Student *t* tests. ***P < .001.

## Discussion

This study identifies coordinated suppression of HNF4α and HNF4γ as a potential mechanism of persistent ileal epithelial barrier dysfunction in CD. Combined loss of both paralogs disrupted transcriptional programs governing TJ assembly, epithelial polarity, junctional-cytoskeletal organization, and junctional repair and produced size-selective paracellular permeability in small-intestinal organoids maintained without inflammatory or stromal cells. HNF4α, HNF4γ, and CLDN15 were coordinately reduced in human ileal CD epithelium, and this HNF4-low phenotype was retained in PDOs derived from macroscopically non-inflamed ileal mucosa. Chronic TNF-driven ileitis recapitulated the phenotype, which was also retained *ex vivo*. NCT and NFT increased HNF4α, HNF4γ, and CLDN15 expression in human CD PDOs, increased HNF4α and CLDN15 in *Tnf*^ΔARE/+^ organoids, and improved epithelial electrical resistance in CD monolayers. Together, these findings define a segment-selective, epithelial-intrinsic, and pharmacologically reversible HNF4-low state that may help explain why ileal barrier dysfunction is detectable in non-inflamed mucosa and can persist despite clinical or endoscopic disease control.^5^

The requirement for combined HNF4α and HNF4γ loss helps reconcile the limited or variable barrier phenotypes reported in HNF4α-only models. Neither single knockout significantly disrupted small intestinal barrier-associated transcriptional programs, whereas combined loss caused broad transcriptional dysregulation and size-selective paracellular permeability. These findings support functional compensation between the paralogs and indicate that HNF4α-deficient models alone do not fully represent HNF4 function in the small intestine. This requirement was strongly segment selective, combined *Hnf4α/γ* deletion extensively disrupted barrier-associated transcriptional programs in the small intestine but altered only 5 TJ-associated genes in the colon. Moreover, *Hnf4α* deletion alone did not significantly alter colonic TJ-associated gene expression, despite HNF4α being the predominant HNF4 paralog in the adult colon.^31, 35^ Because HNF4γ expression is low in the adult colon, this limited response is unlikely to reflect compensation by HNF4γ. Colonic TJ regulation may instead be less dependent on HNF4 activity or maintained by alternative region-specific transcriptional networks. The marked difference between the small-intestinal and colonic responses also argues against the barrier-phenotype being solely a nonspecific consequence of generalized epithelial dysfunction. Although multiple mechanisms likely contribute to regional differences in CD, this segment-selective requirement may provide mechanistic context for the greater prognostic value of terminal ileal compared with colonic barrier function.^5^ The HNF4-dependent small-intestinal defect also appears distinct from acute barrier-disruptive mechanisms commonly described in UC, in which active colonic inflammation promotes cytokine-mediated TJ remodeling, epithelial shedding, and increased paracellular permeability ^8, 58, 59^ Barrier abnormalities have also been reported in clinically quiescent UC, although residual inflammatory activity complicates their interpretation.^60, 61^ Together, these findings identify a segment-selective requirement for combined HNF4-paralog activity in maintaining small-intestinal barrier programs and suggest that this mechanism differs from the predominantly inflammation-associated mechanism in the colon.

The functional studies demonstrate that combined HNF4-paralog loss produces size-selective paracellular permeability. *Hnf4α/γ*^DKO^ organoids accumulated 4- and 10-kDa dextrans while continuing to exclude the 70-kDa tracer, indicating increased permeability without gross loss of epithelial integrity. This phenotype was accompanied by reduced expression of TJ scaffolds, polarity regulators, junctional-cytoskeletal coupling proteins, and mediators of junctional repair. HNF4-paralog loss also increased *Cldn2*, raising the possibility that ion-selective permeability is altered in addition to size-selective dextran permeability.^50, 62, 63^ In a complementary functional assay, CD PDO monolayers exhibited reduced TEER, which increased toward non-CD levels following treatment with NCT or NFT. These findings establish that combined HNF4-paralog loss impairs barrier function in genetically defined mouse organoids and associate the HNF4-low state with reversible barrier dysfunction in human CD-derived epithelium. Neither dextran permeability nor TEER identifies the specific paracellular pathways involved or resolves ion and charge selectivity. Transepithelial conductance, dilution-potential analyses, and matched ion- and size-selective solute flux measurements will be required to define how HNF4-paralog suppression affects distinct components of epithelial barrier function. CLDN15 was therefore examined as a mechanistically anchored readout of HNF4-dependent TJ transcription rather than as the sole mediator of barrier dysfunction. CLDN15 forms a paracellular cation channel that supports Na⁺ recycling and nutrient absorption in the small intestine, and *Cldn15* is a direct HNF4α transcriptional target.^33, 64^ Because restoration of a cation-selective pore would not directly explain increased TEER, recovery of CLDN15 should be interpreted as evidence of broader HNF4-associated transcriptional recovery rather than as the cause of the TEER response.

The segment-selective HNF4 phenotype may be relevant to the progression of ileal CD. Pretreatment ileal transcriptomic studies have identified inflammatory, wound-healing, and extracellular-matrix programs associated with macrophage and fibroblast activation in patients who subsequently developed strictures.^65^ Our findings add a complementary epithelial component to this framework by demonstrating coordinated HNF4α/γ reduction in ileal CD, disruption of HNF4-dependent barrier programs in genetically defined models, and impaired barrier function in CD-derived epithelium. Although progression to fibrostenotic or penetrating disease was not examined directly, these findings support a testable model in which incomplete recovery of the HNF4-dependent epithelial barrier permits continued luminal antigen exposure, thereby sustaining immune and stromal signaling involved in tissue remodeling.^12, 13^ In this model, HNF4 suppression would act as a permissive epithelial defect rather than a direct cause of fibrosis or fistula formation. This hypothesis provides a potential biological link between persistent ileal barrier dysfunction and the greater risk of stricturing or penetrating progression in ileal CD.^5^

Several upstream signals may contribute to the HNF4-low state. Microbiota colonization is associated with a genome-wide reduction in HNF4α and HNF4γ chromatin occupancy in intestinal epithelial cells.^66^ In hepatocytes, inflammatory signaling, endoplasmic reticulum stress, and ERK1/2 activation can suppress HNF4α expression, stability, chromatin binding, or transcriptional activity.^67–70^ Liver regeneration provides a precedent in which transient reduction of HNF4 activity permits proliferative and actue-phase responses, whereas subsequent restoration is required to re-establish differentiated tissue function. ^71–74^ Although these regulatory mechanisms have not been demonstrated directly in intestinal epithelium, they support a model in which transient HNF4 reduction may facilitate an adaptive injury response, whereas failure to restore HNF4 activity contributes to persistent epithelial dysfunction.

The *Tnf*^ΔARE/+^ findings connect chronic inflammatory injury with this model. Coordinated HNF4α/γ reduction was present in chronically inflamed *Tnf*^ΔARE/+^ ileal tissue, and HNF4α, HNF4γ, and CLDN15 remained reduced in organoids after removal from the inflammatory tissue environment. This phenotype paralleled that observed in human CD PDOs derived from macroscopically non-inflamed ileal mucosa. Its retention *ex vivo* indicates that continued exposure to inflammatory, immune, or stromal signals is not required to maintain the HNF4-low state once established. Failure to re-establish the homeostatic HNF4-dependent barrier could help explain why clinical, endoscopic, or histologic improvement does not always result in functional barrier healing.^5^ Prospective studies will also be needed to determine whether ileal HNF4α/γ suppression predicts failure of functional barrier healing or progression to stricturing or penetrating disease independently of inflammatory activity.

Previous efforts to pharmacologically modify epithelial permeability have largely targeted barrier loss occurring during active inflammatory signaling. Blockade of MLCK1 recruitment to the perijunctional actomyosin ring with divertin prevents TNF-induced junctional opening, whereas prednisolone and JAK inhibition protect epithelial and organoid barriers from experimentally impose cytokine injury.^75–77^ EGF and butyrate have also been evaluated as epithelial-supportive therapies, primarily in active colonic disease, although their clinical development has been constrained by safety concerns, inconsistent efficacy, or limited evidence in ileal CD.^78–80^ More recently, TMEM219 blockade restored intestinal stem-cell self-renewal and promoted mucosal healing, but reversal of a permeability defect retained in non-inflamed ileal CD epithelium was not examined.^81^

The present pharmacologic experiments address a distinct epithelial state. CD PDOs exhibited reduced HNF4α, HNF4γ, and CLDN15 expression and impaired barrier function without experimentally added inflammatory cytokines. NCT and NFT increased HNF4α, HNF4γ, and CLDN15 expression and increased TEER toward non-CD levels. Thus, the compounds reversed established molecular and functional abnormalities rather than merely preventing acute cytokine-induced junctional disruption. NCT and NFT also increased HNF4α and CLDN15 expression in *Tnf*^ΔARE/+^ organoids, extending the molecular response to an independent model of chronic ileitis. These parallel responses demonstrate that features of the HNF4-low epithelial state remain pharmacologically reversible across disease-relevant models. *In vivo* studies will be required to define durability, pharmacology, safety, and therapeutic efficacy during chronic ileitis. Nevertheless, these findings support further evaluation of HNF4-directed epithelial restoration as a strategy potentially complementary to control of mucosal inflammation.

### Limitations

The human cohorts were modest in size and the cross-sectional design precluded determining whether HNF4-paralog suppression predicts subsequent progression to stricturing or penetrating disease. Pharmacologic rescue was demonstrated *ex vivo*; the durability, safety, and therapeutic efficacy of HNF4 activation *in vivo* remain to be established.

### Summary

In summary, this study provides a mechanistic explanation for persistent ileal barrier dysfunction in CD that is not accounted for by ongoing inflammatory signaling alone. Combined HNF4α/γ loss disrupted a segment-selective transcriptional program governing TJ assembly, epithelial polarity, junctional-cytoskeletal organization, and repair and was sufficient to produce size-selective paracellular permeability in the absence of inflammatory or stromal cells. HNF4α, HNF4γ, and CLDN15 were coordinately reduced in human ileal CD epithelium and in PDOs derived from macroscopically non-inflamed ileal mucosa. Chronic TNF-driven ileitis recapitulated this HNF4-low phenotype, which was retained in corresponding organoids after removal from the inflammatory tissue environment. These findings support a model in which chronic inflammatory injury establishes or reinforces an epithelial state characterized by sustained HNF4-paralog suppression and incomplete barrier recovery. NCT and NFT reversed the HNF4-low molecular phenotype in CD PDOs and increased epithelial electrical resistance toward non-CD levels. Together, these results identify incomplete restoration of the HNF4-dependent epithelial program as a potential mechanism of persistent ileal barrier dysfunction and support further evaluation of HNF4-directed barrier restoration as a therapeutic strategy complementary to inflammatory control.

## Grant Support

This work was supported by NIDDK (DK-098120, DK-117476 to S.K.; DK-126446 to M.P.V.) with additional support from the Texas Medical Center Digestive Disease Center (P30DK056338).

## Disclosure

The authors disclose no conflicts of interest.

## Abbreviations

CD: Crohn’s disease
CLDN15: claudin-15
DAPI: 4′,6-diamidino-2-phenylindole
DKO: double knockout
DMSO: dimethyl sulfoxide
FITC: fluorescein isothiocyanate
HNF4: hepatocyte nuclear factor 4
IBD: inflammatory bowel disease
KO: knockout
NCT: *N*-trans-caffeoyltyramine
NFT: *N*-trans-feruloyltyramine
PDO: patient-derived organoid
PPARγ: peroxisome proliferator–activated receptor γ
SEM: standard error of the mean
TEER: transepithelial electrical resistance
TJ: tight junction
TNF: tumor necrosis factor
UC: ulcerative colitis
WT: wild type

## Author Contributions

Seema Khurana conceived and supervised the study, obtained funding, contributed to experimental design and data interpretation, and prepared the manuscript. Debasish Halder, Amin Esmaeilniakooshkghazi, Chiranjeevi Padala, Xia Qiu, and Lei Chen performed experiments and analyzed data. Jason K. Hou provided human tissue specimens and contributed to the clinical interpretation of the human Crohn’s disease findings. Michael P. Verzi provided organoids derived from Hnf4α/γ double-knockout and control mice and contributed to the interpretation of the HNF4-paralog findings. Yaohong Wang performed and interpreted the histopathologic analyses. All authors critically reviewed the manuscript and approved the final version.

## Data Availability

The data supporting the findings of this study are available from the corresponding author upon reasonable request. Previously published transcriptomic datasets analyzed in this study are available through the repositories and accession numbers cited in the corresponding publications. Newly generated RNA-sequencing data will be deposited in a public repository before publication.

## Ethical Considerations

All human studies were conducted in accordance with institutional guidelines and approved by the appropriate Institutional Review Board (IRB). Human intestinal tissue samples were obtained with written informed consent and de-identified prior to analysis. All animal experiments were approved by the appropriate Institutional Animal Care and Use Committee and performed in compliance with institutional and national guidelines for the care and use of laboratory animals.

## Notes

### Competing Interest Statement

The authors have declared no competing interest.

### Summary of Updates

We have reanalyzed our RNA-seq data provided in this manuscript and added an additional figure. These changes required significant edits of results and discussion.

